# Network dynamics in the healthy and epileptic developing brain

**DOI:** 10.1101/133488

**Authors:** RE Rosch, T Baldeweg, F Moeller, G Baier

**Affiliations:** Wellcome Trust Centre for Neuroimaging, University College London, UK; Developmental Neurosciences Programme, UCL Great Ormond Street Institute of Child Health, University College London, UK; Department of Clinical Neurophysiology, Great Ormond Street Hospital, London, UK; Cell and Developmental Biology, University College London, London, UK

**Keywords:** EEG, Dynamic Network, Epilepsy, State Transitions, Computational Analysis

## Abstract

Electroencephalography (EEG) allows recording of cortical activity at high temporal resolution. EEG recordings can be summarised along different dimensions using network-level quantitative measures, e.g. channel-to-channel correlation, or band power distributions across channels. These reveal network patterns that unfold over a range of different time scales and can be tracked dynamically.

Here we describe the dynamics of network-state transitions in EEG recordings of spontaneous brain activity in normally developing infants and infants with severe early infantile epileptic encephalopathies (*n*=8, age: 1-8 months). We describe differences in measures of EEG dynamics derived from band power, and correlation-based summaries of network-wide brain activity.

We further show that EEGs from different patient groups and controls can be distinguished based on a small set of the novel quantitative measures introduced here, which describe dynamic network state switching. Quantitative measures related to the smoothness of switching from one correlation pattern to another show the largest differences between groups.

These findings reveal that the early epileptic encephalopathies are associated with characteristic dynamic features at the network level. Quantitative network-based analyses like the one presented here may in future inform the clinical use of quantitative EEG for diagnosis.

## INTRODUCTION

Epilepsy is the most common primary neurological disorder globally, with a particularly high incidence in infancy and childhood (Olafsson et al. 2005). In a group of epilepsy syndromes, the burden of epileptic discharges can cause severe, persistent brain dysfunction, i.e. a recognisable encephalopathy. When these occur in early infancy, they are known as early infantile epileptic encephalopathies (EIEE) (Jette et al. 2015; Ben-Ari & Holmes 2006). Within the category of severe epilepsies, there are several discrete electroclinical syndromes that follow specific developmental timelines, occurring mainly in the neonatal period or very early infancy (e.g. Ohtahara syndrome), later during infancy (e.g. infantile spasms / West syndrome), or in early childhood (e.g. Lennox-Gastaut syndrome). This developmental pattern can also be observed in individual patients, such that a syndromic pattern may evolve e.g. from Ohtahara to West syndrome during development. This suggests that despite an individually persistent cause for the epilepsy (such as a genetic mutation or structural lesion), it is specific stages of brain development that translate the abnormality into age-specific, recognisable electroclinical phenotypes (Kodera et al. 2016; Ohtahara & Yamatogi 2006).

Electroencephalography (EEG) gives a rich picture of dynamic neuronal function and regionally distinct oscillatory brain behaviour in frequency ranges that span several orders of magnitudes (Lopes da Silva 1991). In clinical practice, EEG analysis is focussed on visual pattern recognition of specific waveform abnormalities (e.g. epileptiform discharges) associated with specific clinical correlates (e.g. increased risk of epileptic seizures). Visual analysis – whilst essential – is biased towards certain observable features: For example, between-channel correlation of low frequency, high amplitude discharges is much more readily apparent than of high frequency, low amplitude discharges. Quantitative, automatic analysis may reveal some of these EEG features usually overlooked by visual analysis alone (Tong & Thakor 2009).

Recently graph theory, or network-based approaches to understanding neuronal function in terms of ‘functional networks’ have emerged in imaging neuroscience. Particularly in functional magnetic resonance imaging (fMRI) of the resting state this has led to the discovery that neuronal networks show functionally relevant and quantifiable fluctuation between different constellations, or states over time (Allen et al. 2014; Krienen et al. 2014). Similar methods based on graph theory have now been applied to electrophysiological signals from EEG and MEG recordings in humans (Brookes et al. 2011; Maldjian et al. 2014; Boersma et al. 2011), and suggest that the high temporal resolution in these signals can be harnessed to identify recognisable ‘microstates’ at millisecond-to-second time scales and characterise the switching between them (Van De Ville et al. 2010; Koenig et al. 2002; Baker et al. 2014; Khanna et al. 2015; Vidaurre et al. 2016).

Dynamic features not directly visible in EEG analysis – such as the microstate dynamics described above – are not commonly considered in computational analyses of clinical EEG recordings. There is an emerging literature on the computational analysis of EIEE phenotypes (Japaridze et al. 2013; Japaridze et al. 2016) and related abnormal EEG patterns (Liu & Ching 2017;Ching et al. 2012). Yet our understanding of intrinsic network dynamics in these phenotypes is still limited. Yet, these network dynamics are potentially important to understand whole brain dysfunction as seen in the EIEEs, where there is often not a sharp distinction between seizure patterns and interictal abnormalities.

The work presented in this paper has two main goals: To (1) describe a method to quantify network dynamics in terms of dynamical switches between EEG states based on correlation patterns and power distributions, thus deriving a multivariate feature space capturing network-level brain dynamics. And (2) to evaluate whether this approach captures pathological brain dynamics by mapping two distinct EIEE electroclinical syndromes (Ohtahara syndrome, West syndrome) onto this brain dynamics feature space. In the future, such an approach may prove valuable for resolving diagnostic uncertainties (e.g. in neonatal epilepsy), but also inform computational models of neuronal populations, and thus help identify the neurobiological mechanisms underlying the phenotypes seen in this group of severe epilepsies.

## METHODS

### 2.1 Subjects

This study is focussed on establishing estimates that describe EEG microstate dynamics using different network measures, and thus illustrates the methodology on a small number of participants with profound EEG abnormalities. Both patients and control EEG recordings were selected from previously recorded standard paediatric clinical EEGs. The selection was based on classification by a clinical neurophysiologist with expertise in paediatric EEG (FM).

Subject characteristics are detailed in Table 1. Two control subjects were identified from routine clinical service in a tertiary paediatric hospital providing specialist regional neurophysiology services, based on their age and an EEG within normal limits without evidence of epileptiform abnormalities. Patients with Ohtahara syndrome were selected based on (1) clinical history of seizures, (2) neonatal or early infantile onset of the epilepsy, and (3) evidence of a burst-suppression pattern on standard clinical EEG. Patients with West syndrome were selected based on (1) clinical history of infantile spasms, (2) infantile onset of the epilepsy, and (3) evidence of hypsarrhythmia on standard clinical EEG. Examples of the EEGs from patients, compared to the age-matched healthy controls are shown in Fig 1. Where identified, underlying causes for the epilepsy ranged from genetic abnormalities, localised brain lesions to brain malformations.

**Table 1:** EEG and clinical features of participant included in the analysis.

| ID | EEG Classification | Age (months) | EEG Abnormality |
| --- | --- | --- | --- |
| 1 | Normal for age | 2 | - |
| 2 | Normal for age | 6 | - |
| 3 | Ohtahara Syndrome | 1 | Burst suppression |
| 4 | Ohtahara Syndrome | 2 | Burst suppression |
| 5 | Ohtahara Syndrome | 2 | Burst suppression |
| 6 | West Syndrome | 5 | Hypsarrhythmia |
| 7 | West Syndrome | 6 | Hypsarrhythmia |
| 8 | West Syndrome | 8 | Hypsarrhythmia |

**Figure 1.**
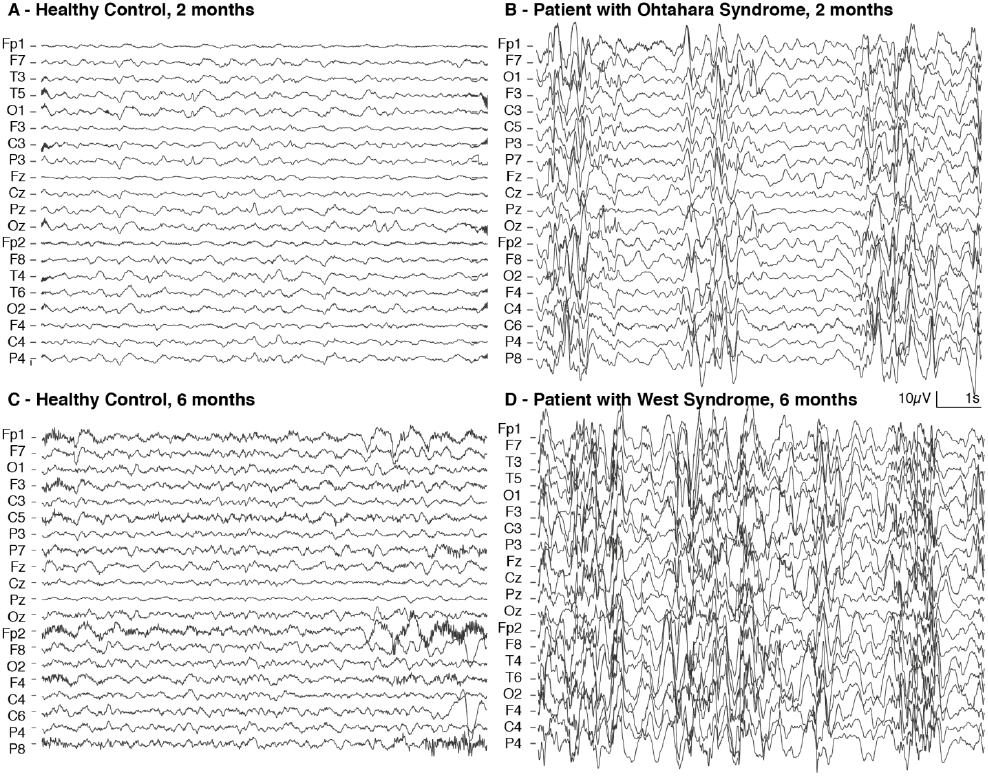
EEG recordings of healthy controls (A,C) and patients with EIEEs (B,D). Compared to healthy controls at 2 months (A), Ohtahara syndrome is associated with abnormal burst-suppression patterns disrupting the ongoing background, characterised by widespread, intermittent bursts of high amplitude activity (B). Compared to healthy controls at 6 months (C), West syndrome is associated with chaotic and disorganised high amplitude activity with mixed frequency components (hypsarrhythmia, D). All EEGs are shown in average referential montage.

All EEG recordings were performed with informed consent from the patients’ legal guardians, and as necessitated by the patients’ clinical course. Use of anonymised EEGs from the clinical database for quantitative analysis was reviewed and approved by the UCL Great Ormond Street Institute of Child Health Joint Research and Development office.

### 2.2 EEG Recordings and Preprocessing

Routine clinical EEG recordings were used for the analysis. Each patient had 19-21 scalp electrodes placed according to the International 10-20 system.

Recordings lasted for up to 30 minutes during task free resting with the subjects’ parent or guardian. Data were recorded with a sampling frequency of 256 or 512Hz and Butterworth bandpass filtered to a 1-80Hz frequency band for visual analysis.

For each individual subject a total of five artefact-free 10s segment of EEG were selected for further analysis, excluding periods of visually apparent deep sleep. No distinction was made for light sleep and awake segments in the EEGs where no obvious electrographic sleep architecture was appreciated on visual analysis. Because of the severity of the EEG abnormalities, sleep stages were not apparent for some of the patients.

Further analysis was performed on these data segments, each filtered to 6 different standard EEG frequency bands: broadband (0.1-60Hz); delta-band (1-4Hz), theta band (4-8Hz), alpha band (8-13Hz), beta band (13-30Hz), and gamma band (30-60Hz). Quantitative analysis was performed on an average montage with each channel referenced to the overall mean scalp activity.

### 2.3 Quantitative Network Analysis

Analysis was performed using customised scripts written by the authors running on <u>Matlab 2016a</u>, as well as the <u>k-Wave toolbox</u> (Treeby et al. 2011) for quantification of matrix sharpness and contrast, and the <u>Chaos toolbox</u> to evaluate stationarity in the features described here. All scripts are available to download and free to use at <u>doi.org/10.5281/zenodo.571647</u>.

#### Estimating dynamic correlation pattern changes

In order to identify changes in network states, dynamic correlation patterns were estimated (summary of the analysis pipeline shown in Fig 2): For each EEG segment a sliding window approach was used to estimate dynamic changes in the patterns of correlation between channels: Starting at each sampling point between 0 and 8s of any 10s EEG segment, a two-second window was extracted, yielding *k* overlapping short segments. For each short segment, pairwise Pearson’s correlation indices between individual channels were calculated, yielding a total of *k* correlation matrices that were *n* * *n* elements large each (Fig 2B, where *k* = number of steps for sliding window, *n* = number of channels).

**Figure 2.**
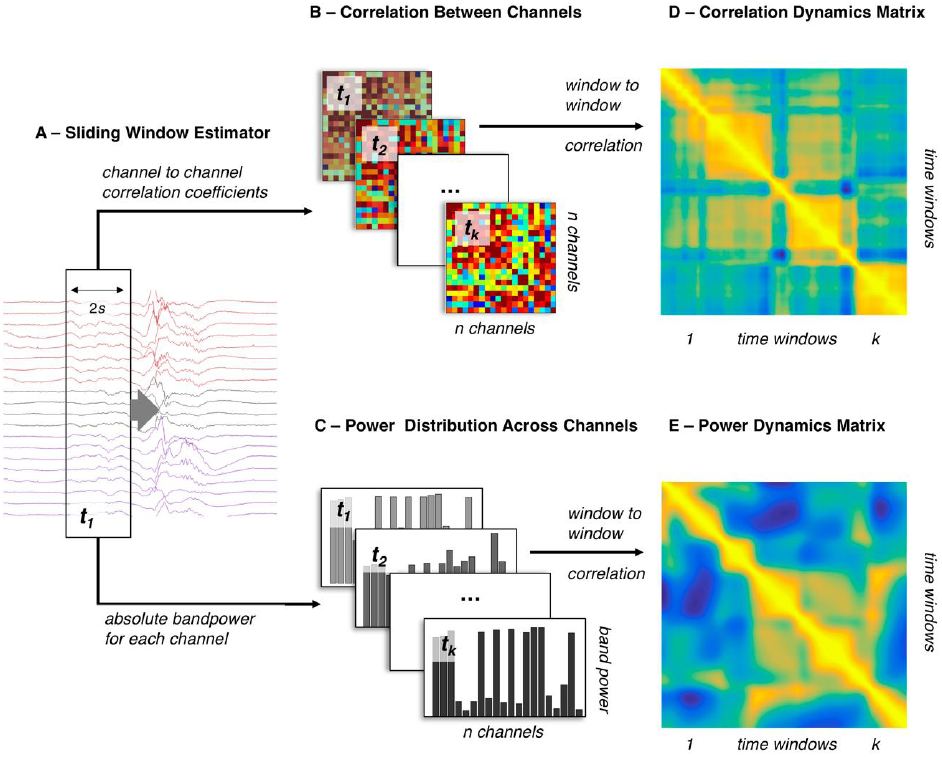
Dynamics of band power fluctuations and changing network correlations can be estimated separately. This figure summarises the estimation of dynamic changes in the power distribution, and correlation patterns over a single 10 second window: (A) A sliding window estimator (window length: 2*s*; step size: 1/512 *s*) is used for feature extraction. (B) Channel-by-channel correlations are estimated using Pearson’s linear correlation coefficients for each time window. (C) The average power within the specified frequency band is estimated for each channel independently, resulting in a specific band power distribution for each time window. (D,E) Estimating window-to-window correlation based on these measures yields two *n* * *n* dynamics matrices describing correlation dynamics (D: CDM), and power dynamics (E: PDM) respectively.

To identify transitions between network states defined by specific scalp-electrode correlation patterns, a single *k* * *k* correlation dynamics matrix (CDM) was calculated to identify temporal changes in the correlation patterns of the channel-to-channel correlation matrices (Fig 2D). Further statistical analysis was based on a single such CDM for each 10s time window.

#### Estimating dynamic power distribution changes

Similar to the correlation dynamics analysis, we established a related measure describing the power distribution changes over time (also shown in Fig 2): For each EEG segment, a sliding window approach was used to calculate mean band power for each channel within the respective frequency band. This yielded *k* vectors of length *n* (Fig 2C, with *k* = number of steps for sliding window, *n* = number of channels). A *k* * *k* power dynamics matrix (PDM) was calculated by estimating the pairwise correlation between each of the *k* power distribution vectors (Fig 2E). Further statistical analysis was based on a single PDM for each 10s time window.

### 2.4 Statistical Analysis

#### Quantifying matrix features

To quantify features in the dynamics matrices, a set of scalar measures was calculated for each dynamics matrix; these are summarised in Table 2. Briefly, they include the matrix mean (i.e. dynamic correlation averaged in time); matrix contrast; and matrix sharpness (defined as the Brenner Operator, Treeby *et al.*, 2011). Contrast and sharpness measures are derived from image analyses and in this context, represent measures describing the transition between correlated network states in time. They are differentially sensitive to regional amplitude differences (where contrast is more robustly sensitive) and smoothness of transition between states (where sharpness is more robustly sensitive), as illustrated in Fig 3.

**Table 2:** Measures used for quantification of dynamics matrix features

|  |  |
| --- | --- |
| Mean values | $\sum_{x,y} f_{x,y} / k^2$ |
| Contrast* | $\sum_{x,y} (f_{x,y} - f_{x+1,y+1})^2$ |
| Sharpness (Brenner Operator)** | $\sum_{x,y} (f_{x+2,y} - f_{x,y})^2 + (f_{x,y+2} - f_{x,y})^2$ |

\* derived indirectly from the gray level co-occurrence matrix
\*\*modified from Treeby *et al.*, 2011
$f_{x,y} = f(x, y)$ , the scalar value of the matrix at column $x$ , row $y$ . $k$ is the number of rows in the matrix

**Figure 3:**
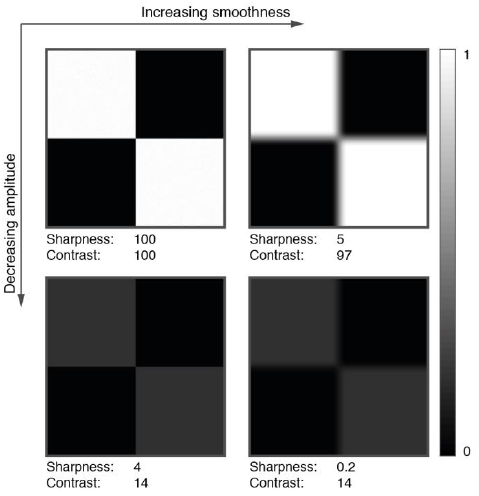
Contrast and sharpness measures encode related but different matrix features. Analysis of example matrices with different levels of Gaussian smoothing (left vs right panels), and different signal amplitudes (top vs bottom panels) shows differential sensitivity of the measures employed (normalised values shown): Contrast changes mainly with amplitude of the regional signal differences, whilst sharpness is affected both by amplitude, and smoothness modulations.

#### Testing for stationarity and statistical differences

To evaluate whether the dynamic matrix features reflect non-stationary processes,they were statistically evaluated against a set of stationary surrogate time series of the same frequency composition. For each individual time window included in the analysis, the following analysis steps were performed in order to derive a normal distribution of the measures illustrated above (mean values, contrast, and sharpness) in a stationary time series the following steps were performed:

1. Fourier transforms for each channel were calculated for the whole length of each 10s EEG segment
2. A total of 50 amplitude adjusted surrogate time series were generated for each channel based on their Fourier transforms (based on Schreiber and Schmitz, 1996, using the <u>Chaotic Systems Toolbox</u>)
3. Each set of synthetic time series were then analysed using the same sliding window analysis applied to the original datasets, thus deriving 50 surrogate sets of the measures described above for each 10s time window analysed.
4. z-scores were calculated for each measurement derived from the empirical time series, based on the distribution of synthetically generated surrogate measurement sets

If a dynamics matrix measure reflects only stationary processes caused by random fluctuations around steady-state frequency distributions, empirical values are expected to fall within the normal distribution of the surrogate data, i.e. roughly within the −2 to 2 z-score interval. We also used the z-score normalised data to test for differences between individual measures using two-sided *t*-tests.

#### Clustering based on network dynamics

The approach taken above yields several distinct measures for each EEG time window: 3 dynamics matrix measures (mean, contrast, sharpness) for 2 different matrix types (CDM, PDM), for 6 frequency bands (broadband, delta, theta, alpha, beta, gamma) expressed as 2 values (raw, z-score) results in a total of 72 feature measures for each EEG window. To identify dynamic network measures that capture discriminatory features between the EEG abnormalities analysed here, we (1) measured according to how well they can be used to classify individual EEG segments into distinct groups, and (2) used a subset of the highest-ranking measures to automatically identify clusters within the data using machine learning approaches.

For each measure we identified thresholds that optimise the classification of EEG windows into three clusters corresponding to the three participant groups (i.e. Ohtahara syndrome, West syndrome, and healthy controls). To that effect, the purity of the classification, *P*, resulting from a set of two threshold parameters was maximised using a simulated annealing approach. *P* ranges from 0 (no element is correctly categorised) to 1 (all elements are correctly categories) and is calculated as follows:

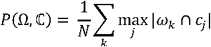

Where Ω= {*ω*, *ω*,…,*ω*}isthe set estimated clusters, and ℂ = = {*c*_1_, *c*_2_, …, *c*_*j*_} is the set of given classes

The maximally achieved value for *P* was recorded for each measure and used to rank the 72 individual dynamics measures according to how well they can be used to cluster EEGs into patient groups. As a second step, we then used subsets of the high-ranking dynamics measures to automatically cluster the EEGs into different groups using k-means clustering. This approach partitions a dataset into a set of *k* clusters automatically, given a set of observations. We use this approach to quantify how well the measures we identified in the first steps can be used to categorise 10s segments of EEG into the appropriate participant categories (using the purity measure *P*), and how well they distinguish between normal and abnormal (Ohtahara, and West syndrome combined) categories (using sensitivity and specificity estimates). This does not aim to assess individual measures in terms of their diagnostic accuracy, but quantify how much information about the original classification based on full EEG recordings is retained in this low-dimensional feature space.

## RESULTS

### 3.1 Correlation and band power dynamics

A total of 5 relatively artefact free EEG segments were selected randomly and analysed for each participant, yielding separate CDM and PDM for each segment and each frequency band. Across all subjects and segments there are visible differences in the temporal dynamics of correlation patterns and band power distribution patterns (shown in Fig 4A-B, which can be quantified in the difference between the two matrices: if band power distribution and channel-to-channel correlation followed the same dynamics, the differences would be expected to center around 0. The mean of the CDM - PDM difference is shown for each time window in the analysis in Fig 4C. This suggests that in healthy controls, PDM values are overall higher than CDM values, which is also seen in Fig 4B. In both patient groups, there is more temporal-cross correlation in network correlation states, than in the band power distribution, resulting in positive mean difference values.

**Figure 4.**
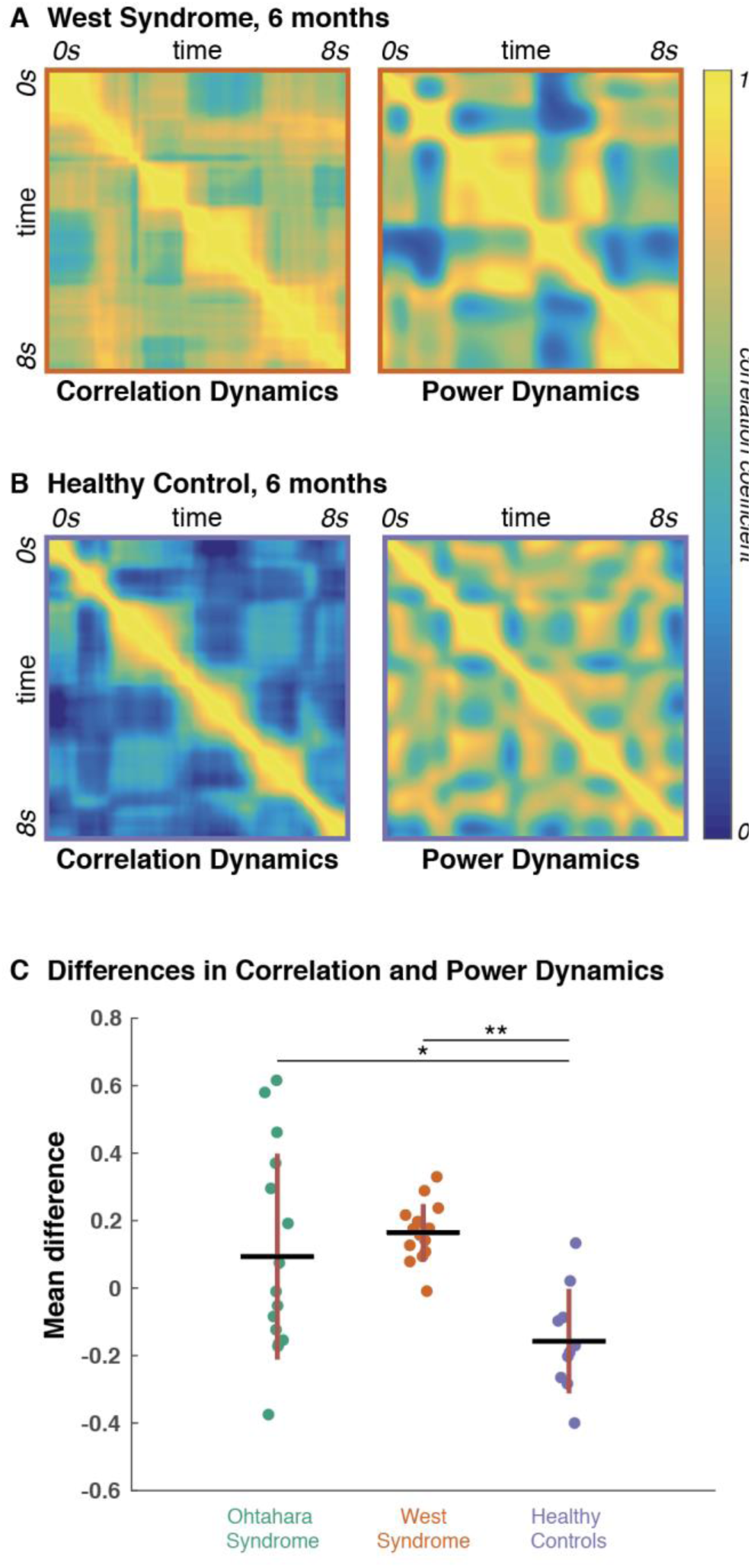
Network correlation states and band power distribution show different recurrence patterns in time. The dynamics matrices show recurrent correlation or band power distribution patterns in time. (A) Shows CDM and PDM for a single 10s EEG segment of a patient with West syndrome. Both show high correlation values outside of the leading diagonal (i.e. between different time segments). (B) In healthy controls, high between time-window correlation is largely restricted to the leading diagonal in the CDM, but not the PDM, suggesting that network correlation patterns are less recurrent than band power distribution patterns. (C) Mean CDM - PDM difference values suggest that for healthy controls, but not the patient groups, recurrent band power patterns recur more across time than network correlation patterns.

A closer analysis of the temporal patterns underlying these differences is shown for a single healthy control EEG segment in Fig 5: Transitions between network motifs as measured through CDM, or PDM show a dissociation: Observed band power distribution across the scalp may change without closely associated corresponding changes in the network correlation patterns and vice versa.

**Figure 5.**
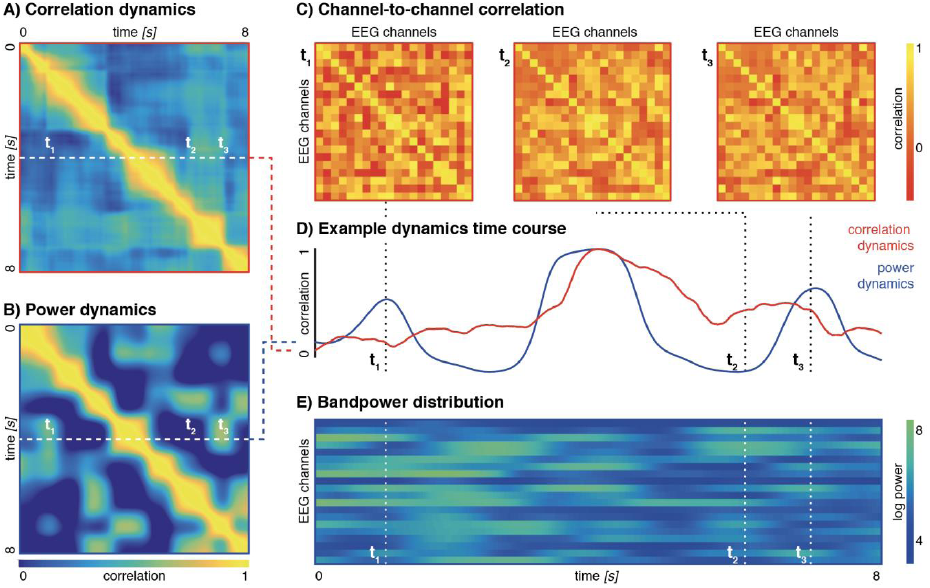
Dissociation of bandpower and network correlation dynamics. (A) Shows the CDM for a single 10s EEG broadband segment in healthy control aged 2 months. (B) Shows the PDM for the same EEG segment. (C) Channel-to-channel correlation patterns are shown for three separate time window, with big differences between time point 1 and the others, and more similarities between time points 2 and 3. (D) Correlations are shown between a single time window (indicated by the dashed lines in A,B) and all other time windows based on correlation dynamics (i.e. CDM) and power dynamics (i.e. PDM). (E) Shows the bandpower distribution across channels over time. Whilst there are broad similarities in the temporal trajectories of power and correlation dynamics, there are discrete instances where similar bandpower distribution patterns (at t_1_ and t_3_) are associated with very different correlation patterns, and vice versa (at t_2_ and t_3_).

### 3.2 Non-stationarity in CDMs

Randomly generated stationary time series of the same spectral composition as the empirical recordings were used to assess for non-stationarity in different measures applied to the EEG segments. For each measure, z-scores were calculated from a distribution generated based on analysis of 50 synthetic data sets for each individual EEG segments, and are shown in Fig 6. Most of the dynamics measures derived from the PDM can be explained as random fluctuations around a stationary distribution, whilst CDM-derived measures differ significantly from the stationary distributions, i.e. show non-stationarity.

**Figure 6.**
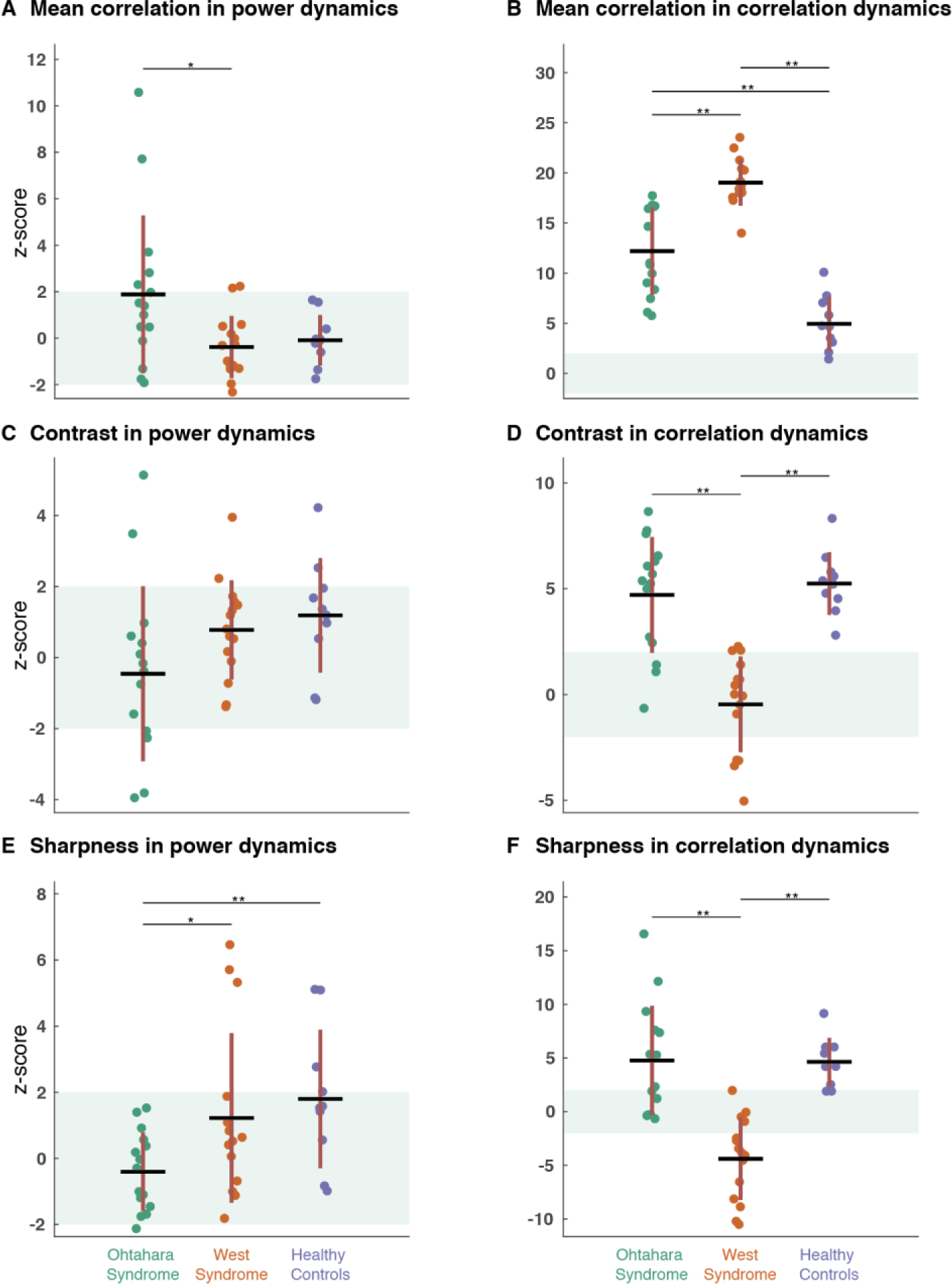
Correlation dynamics are non-stationary. PDM-derived measures of band power dynamics (A,C,E) fall mostly within a 95% confidence interval derived from stationary surrogate synthetic datasets (shown as green shading), indicating that they may represent stationary processes. CDM-derived measures (B,D,F) are not fully explained by random correlations in stationary signals and show significant group differences even when z-score normalised. Red bars indicate the standard deviation around the means of the individual groups.

All z-score normalised CDM measurements show group differences between patients with Ohtahara, or West syndrome and healthy controls. Example CDMs for each participant group are shown in Fig 7A. It is worth noting that direction of the statistical differences is not necessarily evident from the CDM alone: For example, CDM contrast in patients with West syndrome is lower than expected, even though the example CDM for West syndrome shown in Fig 7A appears rich in contrast. This illustrates that expected ranges of CDM measures are highly dependent on the overall frequency composition of the underlying EEG signal, i.e. higher correlation values may be occur by chance if the signal contains a large low frequency component.

**Figure 7.**
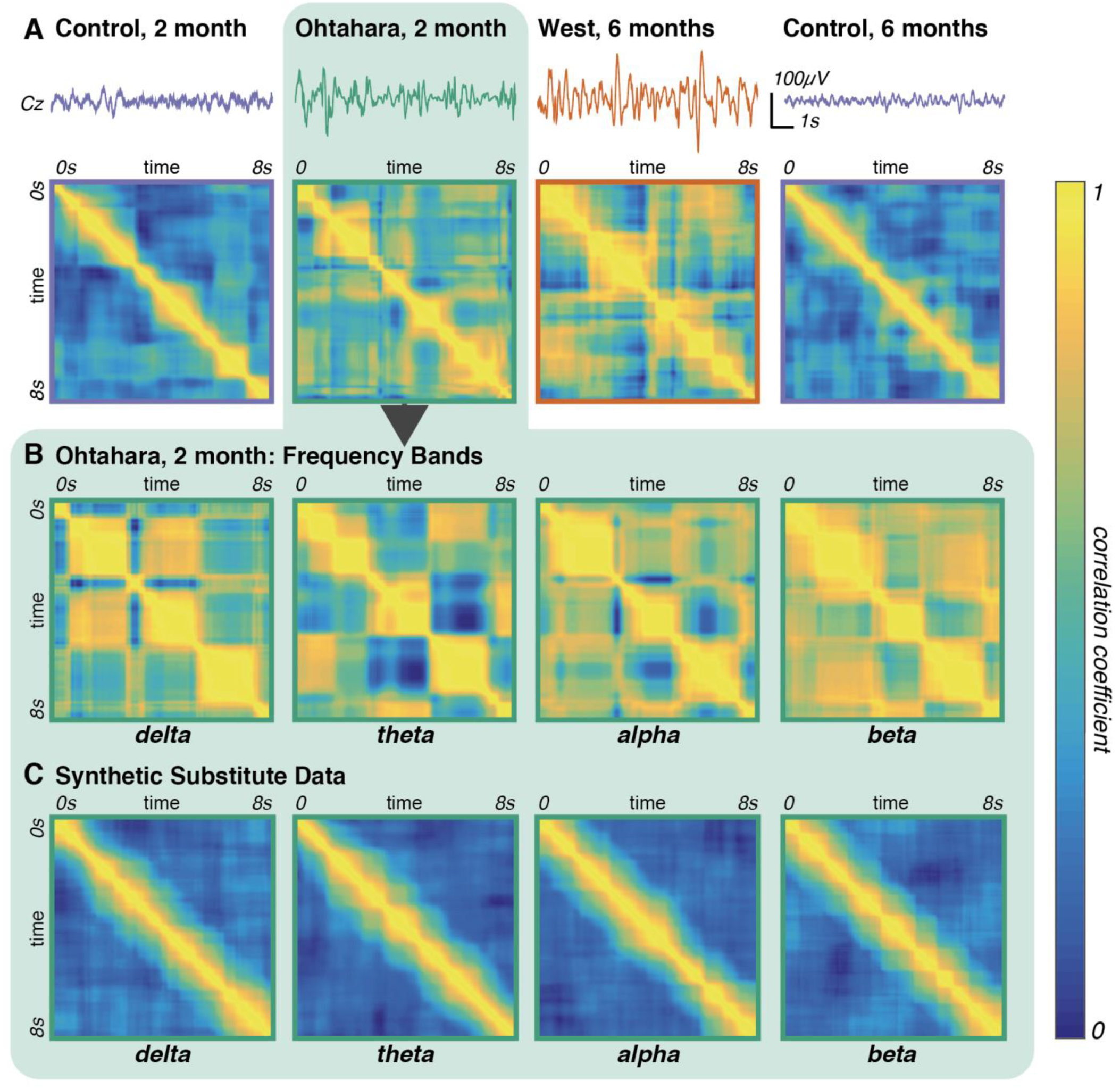
Recurrent and persistent motifs of network correlation states are more apparent in both patient groups. (A) Broadband Cz-electrode time series, and CDMs for are shown for single 10s EEG segments for representatives of both Ohtahara and West syndrome cohorts and healthy age matched controls. (B) CDMs derived from bandpass filtered data for the EEG segment from the Ohtahara patient show distinct dynamics patterns for each frequency band. (C) CDMs derived from stationary surrogate data are largely restricted to the leading diagonal, thus indicating few persistent or recurrent network patterns in any of the frequency bands

### 3.3 Network dynamics in EIEE patients

A complete set of dynamics measures derived from both CDM and PDM were collated for each EEG segment resulting in 72 measures (3 measures * 2 matrix types * 6 filter bands * 2 normalisation types) per EEG segment. For each individual value thresholds were identified that could be used to separate the data into three clusters that reproduced the original participant groups (Ohtahara syndrome, West syndrome, and healthy controls) most closely. Table 3 shows the ranking based on maximum purity achieved for each measure after automatic threshold optimisation. Several of these variables can be used in conjunction to map out distinct groups’ distributions in a low-dimensional feature space. As an example, Fig 8 shows each individual EEG segment mapped onto the three highest ranked dynamics measures from Table 3. In order to verify how well these measures separate distinct subgroups, we evaluated results from a k-means automatic clustering algorithm based on increasingly large subsets (range: 1 to 30 parameters) of the dynamics measures. Results of this unsupervised clustering approach were then compared with the known disease category, using overall classification purity, as well as sensitivity and specificity of separating healthy controls segments from patients (Fig 8B). An example of such a clustering is shown for just two parameters in Fig 8C. Using the top-ranking five parameters, this approach can reach a classification purity of 82.5%, with a disease classification sensitivity of 93.3% and a specificity of 80.0% (Fig 8B).

**Table 3:**
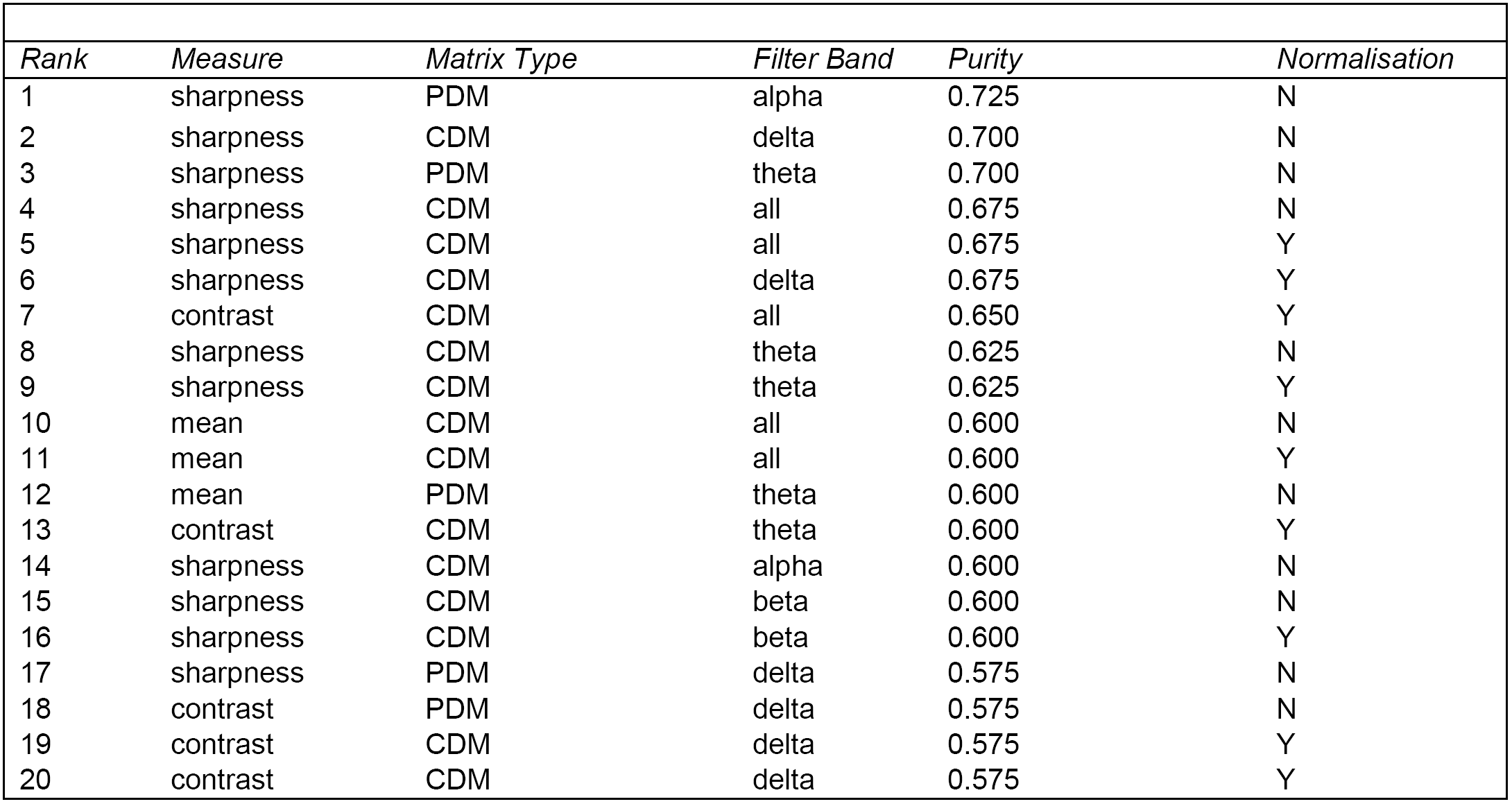
Rank of dynamics measures based on clustering ability

**Fig 8:**
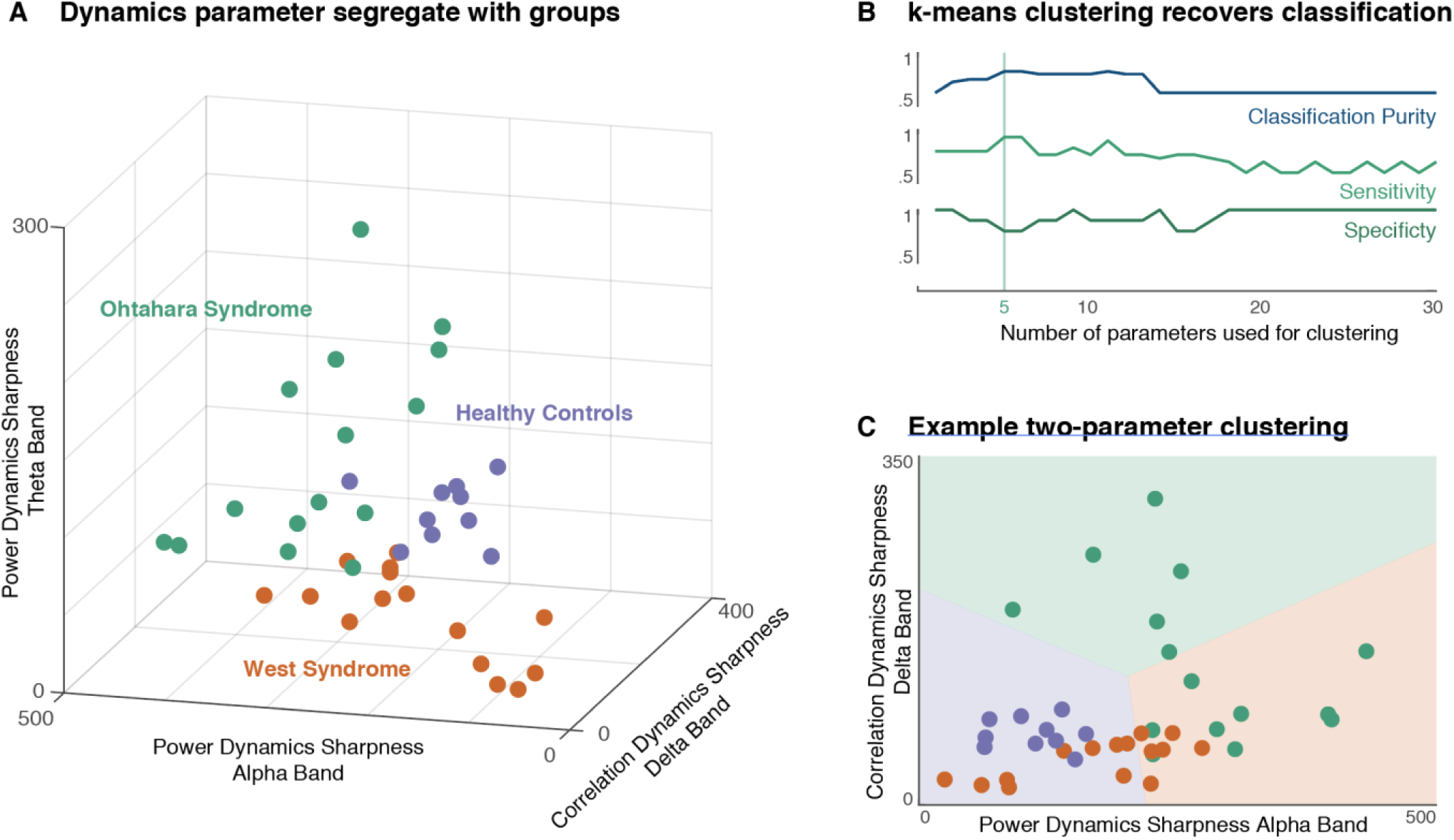
Clustering using dynamics measures separates patient groups. (A) Dynamics matrix measures can be used to visualise clustering of the EEG segments in a three-dimensional feature space. Here all EEG segments are mapped onto the three most discriminatory dynamics measures, producing distinct clusters in this three-dimensional feature space. (B) Classification purity peaks when between 5-15 parameters are used for classification, after which no further gains are made. Sensitivity and specificity of classifying patients and healthy controls in separate groups is also shown. (C) One example clustering solution is shown using only the two highest ranking parameters with limited separation between West syndrome and healthy control EEGs

## DISCUSSION

This report presents a quantitative analysis approach for identifying temporal patterns in network states in the developing brain. Using electrographically distinct epilepsy syndromes affecting most of the background EEG dynamics as illustrative cases we show that (1) temporal correlation analysis can reveal distinct patterns from high dimensional datasets such as EEG; (2) band power and channel-to-channel correlation dynamics can be dissociated, even in the healthy brain; (3) quantitative summary measures derived from this analysis can capture EEG differences between different electroclinical syndromes.

The novel measures introduced here describe quantitatively the temporal dynamics of whole brain network states. Even structurally very different networks may show similarities in dynamics, which makes the measures introduced here particularly useful for identifying similarities in highly heterogeneous clinical populations. Early infantile EEGs are furthermore characterised by activity in a variety of spatial distributions (rather than the more typical posterior dominant rhythms seen in the mature EEG), indicating that analysis of the dynamics of different network patterns unfolding may be particularly informative in this age group.

### Dynamics matrices can reveal hidden temporal structure in high-dimensional data

Network based analyses have been the conceptual basis for the recent success in resting state fMRI in humans (Van Dijk et al. 2010). Furthermore, computational modelling approaches have enabled an understanding of the relationship of observable, macroscopic whole-brain network dynamics and local, mesoscale neuronal dynamics (Deco et al. 2011). Many of the network features first described based on resting state fMRI are also present in the analysis of EEG/MEG recordings, where they can be measured with very high temporal resolution, revealing fast, sub-second recurrent network switching (Baker et al. 2014). These fast network dynamics can be task related (O’Neill et al. 2017), and approaches similar to the one presented here have demonstrated that they are modulated by cognitive tasks, even in children (Dimitriadis et al. 2015).

The analysis presented here is focussed on identifying quantitative EEG features that (i) can show differences between pathological and healthy brain dynamics even at the level of individual subjects, and (ii) can be applied to task-free resting state EEG recordings routinely performed in a clinical setting. Given the heterogeneities in the clinical sample, the aim is not to identify specific neuroanatomical networks that reproduce between these patients, but describe the visually apparent dynamic features using novel quantitative measures and identify whether other, less directly visible features can also be useful discriminators.

The dynamics matrices reveal structured patterns that capture the recurrence of correlated network states over different time scales. The dynamics of these transitions can be visually represented and, importantly, quantified using image metrics (such as sharpness, and contrast) that intuitively capture features of the microstate transitions. This approach summarises specific aspects of the multi-channel, highly time resolved EEG recordings that are less amenable to standard visual analysis.

### Band power and correlation patterns represent different aspects of neuronal circuitry function

Correlations between channels, and power distribution across channels are likely to represent physiologically separate processes that can follow distinct dynamic patterns: The spectral composition of the EEG signal (and thus its regionally specific band power distribution) is believed to result from synchronous firing within local neuronal populations; statistical correlation between distant channels is believed to be caused by direct or indirect, long-range synaptic connectivity (Buzsáki et al. 2012; Vanhatalo & Kaila 2006). A difference between power- and correlation-derived network dynamics can therefore be understood to represent separately the dynamics of synaptic connectivity at the local, and the network level.

Across all participant groups, there are visible differences between dynamic patterns as derived from band power distributions (PDM) and network correlation patterns (CDM). These differences suggest that a particular band power distribution across the scalp do not correspond to network correlation patterns – i.e. functional connectivity motifs – in the developing brain, both in patients and in normally developing controls. Using the approach shown here, changes in correlation patterns and band power patterns over time can be tracked separately and reveal divergent trajectories. Most of the dynamic variance contained within the PDM can be explained as random fluctuations of an overall stationary process, whilst correlation patterns over time differ significantly from those observed in a stationary signal, thus the product of a non-stationary process.

The first year of life is associated with a range of developmental changes affecting both local microcircuitry (e.g. synaptic pruning, neurotransmitter and -receptor changes), as well as global network integration (e.g. myelinisation of large white matter tracts) (Dehaene-Lambertz & Spelke 2015). Identifying developmental changes in the dynamics at different neuronal scales may thus provide insight into the relationship between neurobiological changes and observed EEG patterns, particularly where there are visible EEG changes, as is the case in the EIEEs discussed here. Applied to a larger cohort, the approach illustrated here with just a small sample of pathological EEG patterns may in future also reveal more subtle developmental patterns in the healthy developing brain by allowing quantification of network dynamic behaviours in simple network-based measures, such as the transition sharpness included in the analysis here.

### Quantifying abnormal brain states in clinical populations

Even as little as five scalar measures derived from the network dynamics analysis here can be used to classify EEG segments with reasonable accuracy into the known disease categories. Both Ohtahara and West syndrome are characterised by pervasive neuronal abnormalities that disrupt normal background EEG function. Their associated EEG phenotypes (i.e. burst suppression patterns, and hypsarrhythmia) are readily apparent throughout most EEG segments. Thus, the analysis approach presented here is not designed to resolve diagnostic uncertainty, but the distinct phenotypes included in the analysis were utilised to test whether our novel dynamics measures can reveal apparent differences in the EEG dynamics quantitatively.

As the measures derived from the dynamics matrices are quantifiable, they allow for statistical testing and the application of simple machine learning tools. As illustrated in the approach taken here, individual measures can be ranked according to their discriminative power for clustering into disease categories, thereby identifying the features that most help differentiate different pathologies from the dynamics in the normal developing brain.

Of the ten measures that are most distinct between different groups, eight are measures of sharpness in either the CDMs or PDMs – thus different patients groups and healthy controls show particular differences in their transition between different network states. The majority of the useful measures are derived from the CSMs, suggesting that it is specifically the temporal dynamics of the switch between discrete functional connectivity patterns that separates the groups. Notably, each of the top ten ranking measures were either broadband measures, or restricted to the lower frequencies (delta, theta, alpha), which may be related to the window length of 2s, as this will average out changes in high frequency correlation patterns that are only a few cycles long. However, most of the physiological and abnormal activity we were aiming to capture is within the lower frequency ranges, which the window length appears to capture well.

For almost all measures, Ohtahara syndrome shows a higher sharpness value, suggestive of more acute transitions between more discretely defined states (which may not always correspond to apparent burst activity). Somewhat surprisingly, healthy controls are often found at intermediate values, with West syndrome patients with the lowest sharpness values (e.g. Fig 8A). Yet at the same time both patients with Ohtahara syndrome and with West syndrome show abnormally high persistence of network states (as measured by the mean correlation over time in the CDM, Fig 6B). These observations suggest that the EIEE brain is susceptible to enter recurrent and abnormally stable functional connectivity states, but the expression of dynamic transitions in and out of these is specific to the electroclinical syndrome (and therefore the developmental stage).

This approach complements existing computational modelling of epilepsy and seizure activity. The application of dynamic systems mathematical approaches to neuronal oscillators has led to the recognition of certain stereotypical seizure patterns as mathematically predictable oscillatory patterns (Izkhikevich 2000) that can be reproduced in computational simulations (Jirsa et al. 2014). Using these *in silico* simulations means that we can explain the effects of genetic mutations (Peters et al. 2016), specific seizure responses to stimulation (Taylor et al. 2014), or seizure spread patterns (Baier et al. 2012) using full generative models that bridge observable and non-observable (‘hidden’) spatial and temporal scales. Such approaches focus on reproducing specific features observed in empirical data – typically the particular, directly visible oscillatory patterns and their relationship to neuronal function (Jirsa et al. 2014; Breakspear 2005). Here we offer quantitative descriptions of network level dynamic features that appear to be modulated both by the epilepsy (i.e. persistent network states as seen in the examples in Fig 7A, resulting in the differences in mean correlation apparent in Fig 6B) and developmental stage (i.e. transition dynamics differences between the younger Ohtahara syndrome, and the older West syndrome cohort as indicated in Fig 6D and F) – features that can be specifically included in future models of EIEE.

### Future applications in the clinic and in epilepsy research

Firstly, the EEG phenomenology-based clustering approach may aid in resolving diagnostic uncertainties in neonatal EEG analysis, where the difference between abnormal patterns and normal developmental variants are more difficult to identify visually (Torres & Anderson 1985; Stevenson et al. 2015). Correct diagnosis currently relies on the expertise of clinicians trained in paediatric (and specifically neonatal) clinical neurophysiology, who are not available in all clinical settings where accurate diagnosis of neonatal EEG patterns could be valuable. Yet there is a recent focus on improving neurological outcomes of neonatal care, which is likely to involve a significant increase in EEG recordings and monitoring in neonates at risk of seizures, requiring a corresponding scaling up of EEG interpretation capacities (Vesoulis et al. 2014; Sands & McDonough 2016). Utilising quantitative, computational approaches as presented here may be able to support correct diagnosis in those settings and play a role in improving clinical outcomes (Mathieson et al. 2016; Temko et al. 2011).

Secondly, the epilepsy syndromes under investigation here have a close relationship to developmental stages: Ohtahara syndrome typically is restricted to the neonatal or early infantile period, whilst West syndrome emerges in later infancy, typically between 3 and 10 months. Both share genetic causes (e.g. *GABRA1* mutations, Kodera et al. 2016) and individual patients can evolve from Ohtahara syndrome to West syndrome during their development (Ohtahara & Yamatogi 2006). Thus understanding the neurobiological processes underlying the EEG phenotypes offers a window into the interactions between brain development and early onset pathological processes. By being able to quantify differences in network dynamics we can identify features that are crucial in distinguishing patients groups. These quantitative features can be used as benchmarks for adapting existing models of neuronal dynamics (Baier et al. 2012; Proix et al. 2014; Papadopoulou et al. 2015) to reproduce the empirical observations. With those models we will be able to test mechanistic hypotheses that link recent discoveries on the genetic basis of many of the EIEEs and our understanding of developmental processes in the infant brain, to the identifiable EEG syndromes seen in patients.

### Limitations

This study is not an attempt at testing the dynamics measures in terms of their clinical validity: The EIEE syndromes included here were deliberately chosen because of their wide-ranging impact on the background EEG and the disruption of normal brain dynamics; they are used to illustrate the validity of the method and the possibility to identify less visible group differences. Visually observed EEG differences are large, thus we have only included a small number of subjects, aiming to identify group differences with large effect sizes that are likely to be useful in future applications in clinical samples where predictive power at the single individual is required.

## Acknowledgements

RER is funded by a Wellcome Trust Clinical Research Fellowship (106556/Z/14/Z).

